# Biotic interactions shape infection outcomes in *Arabidopsis*

**DOI:** 10.1101/2024.10.25.620230

**Authors:** Maryam Mahmoudi, Juliana Almario, Yiheng Hu, Lynn-Marie Tenzer, Kay Nieselt, Eric Kemen

**Affiliations:** Microbial Interactions in Plant Ecosystems, IMIT/ZMBP, Eberhard Karls University of Tübingen, Auf der Morgenstelle 32, 72076 Tübingen, Germany; Université Claude Bernard Lyon 1, Laboratoire d’Ecologie Microbienne, UMR CNRS 5557, UMR INRAE 1418, VetAgro Sup, 69622, Villeurbanne, France; Institute for Bioinformatics and Medical Informatics, Eberhard Karls University of Tübingen, Sand 14, 72076 Tübingen, Germany

**Author notes:** **Corresponding author:** Eric Kemen, Microbial Interactions in Plant Ecosystems, IMIT/ZMBP, Eberhard Karls University of Tübingen, Auf der Morgenstelle 32, 72076 Tübingen, Germany.

**Keywords:** Microbe-microbe interaction, infected and uninfected leaves, machine learning, plant pathogen, natural probiotic

## Abstract

The plant microbiome protects plants from stresses, including pathogen attacks. However, identifying microbes that provide plant protection remains challenging in complex microbial communities. In this study, we analysed samples from natural *A. thaliana* populations, including both plants infected with the pathogenic oomycete *Albugo laibachii* and uninfected plants, over six years. Using machine learning classification models, we achieved high accuracy in distinguishing infected and uninfected plants based on microbiome abundance. We identified 80 key taxa associated with health and disease. Among the health-associated microbes (HCom), we selected bacteria, fungi, and cercozoa that effectively reduced pathogen presence in co-inoculation assays. In comparison, disease-associated microbes (DCom) were less effective in conferring protection. Our findings highlight the complexity of plant-microbe interactions and advance our understanding of microbial roles in plant disease ecology. By integrating ecological insights with machine learning, we take a significant step towards designing robust microbial consortia that enhance plant resilience against pathogens.

## Introduction

Similar to higher organisms such as humans, plant tissues are colonized by a wide range of microbes known as microbiota or microbiome. The microbiome associated with the plant leaves, i.e., the phyllosphere, is thought to play an important role in the physiology, fitness, and defense mechanism of the host against various biotic and abiotic perturbations [1]. Leaves are inhabited mainly by commensal species but can also harbor pathogenic bacteria, fungi, and oomycetes, which can significantly harm natural plants and crops [2], thereby causing annual crop yield losses and reducing food availability. Climate change is accelerating the spread of pathogens, thereby affecting forest health globally [3, 4]. Therefore, it is crucial to develop strategies that protect from pathogens in a changing environment.

Biological control is an effective and environmentally friendly alternative to pesticides for combating microbial plant diseases [5]. For example, *Trichoderma*, an opportunistic fungal genus, is widely used as a biological control agent against phytopathogens and is studied for its role in helping plants manage biotic and abiotic stresses [6]. However, identifying and experimentally validating biocontrol microbes through traditional methods can be slow and challenging [7]. Another approach to biocontrol involves identifying healthy microbiomes. While healthy plant communities are characterized by diverse and balanced microorganisms, as seen in comparisons between healthy and diseased plants [8]. However, it remains unclear what defines a healthy or beneficial microbiome. One promising solution to this challenge is the use of machine learning. Machine learning techniques have been used in microbiome research to accomplish various tasks, such as predicting host or environmental phenotypes and categorizing microbial properties, including monitoring changes in microbiome composition [9]. Machine learning classifiers were used to identify soil microbial patterns predicting the presence of *Fusarium oxysporum*, the pathogen causing Fusarium wilt disease under field conditions [10]. Similarly, the random forest method accurately predicted productivity based on microbiome composition at the order level. Significant differences in crop yield were associated with bulk soil microbiome composition, with many taxa contributing to nitrogen utilization [11]. Machine learning was used to identify bacterial strains important in reducing leaf infection with the pathogenic bacterium *Pseudomonas syringae* DC3000 [12]. However, few studies have investigated the different taxonomic groups of bacteria and eukaryotes in the microbiota of natural plants attacked by obligate biotrophic pathogens.

The obligate biotrophic oomycete *Albugo laibachii* is a common pathogen of the *Brassicaceae* family and the causal agent of the white rust disease [13]. This pathogen was identified as a potential core and hub microbe in the leaf microbiome of *A. thaliana* since it showed persistence over several years and high interconnection in the microbial interaction network [14, 15]. *Albugo* infection was also shown to affect both epiphytic and endophytic bacterial colonization by reducing alpha diversity and secretion of antimicrobial peptides [15, 16]. However, it is not clear how the microbiome of the leaf differs in plants infected or not with *Albugo*, and which microbial strains have the potential to promote or reduce infection with *Albugo*.

In this study, we analysed the microbiome of *Arabidopsis thaliana* over a period of six years as collected and described by Mahmoudi et al. [17]. High-throughput sequencing analysis revealed differences in the microbiota composition associated with *Albugo*-infected and uninfected plants across host genotypes and sampling sites. Using statistical and machine learning classification algorithms, we identified candidate microbes predictive of infected and uninfected states. Candidate microbes were shown to be distributed in different clusters in microbial interaction networks, highlighting their importance in the community’s stability. Co-inoculation assays in *A. thaliana* confirmed the potential of the health-associated microbial communities (HCom) to reduce *Albugo* infection. In comparison, disease-associated microbial communities (DCom) exhibited a range of functions, from minimal effects to partial pathogen suppression, likely through niche competition. These findings highlight the functional redundancy of microbial communities from different phylogenetic groups in manipulating plant health outcomes and demonstrate the power of machine learning in informing biocontrol strategies.

## Results

### Comparison of phyllosphere microbiome in natural *A. thaliana* populations: uninfected vs. infected with the obligate biotrophic oomycete pathogen *Albugo*

To investigate the diversity and compositional dynamics of the phyllosphere microbiome in the presence of the obligate biotrophic oomycete pathogen *Albugo*, we used a microbiome dataset described in Mahmoudi et al. [17]. In their study, *A. thaliana* samples were collected from six sites near Tübingen (southern Germany) with stable *A. thaliana* populations, with sampling repeated over six consecutive years (2014-2019). Genomic DNA was extracted from epiphytic and endophytic microbial communities, followed by amplicon sequencing for bacterial 16S rRNA, fungal ITS2, and eukaryotic 18S rRNA. For the 18S eukaryotic data, fungal microbes were excluded, resulting in the nonfungal eukaryotes (NFEuk) dataset [17]. Here, we used the endophytic microbiomes for further analysis. Since *Albugo* was the major pathogen associated with *A. thaliana* at the time of sampling, the samples were categorized as infected or uninfected based on the presence or absence of white rust on the leaves (Fig. 1A).

**Figure 1.**
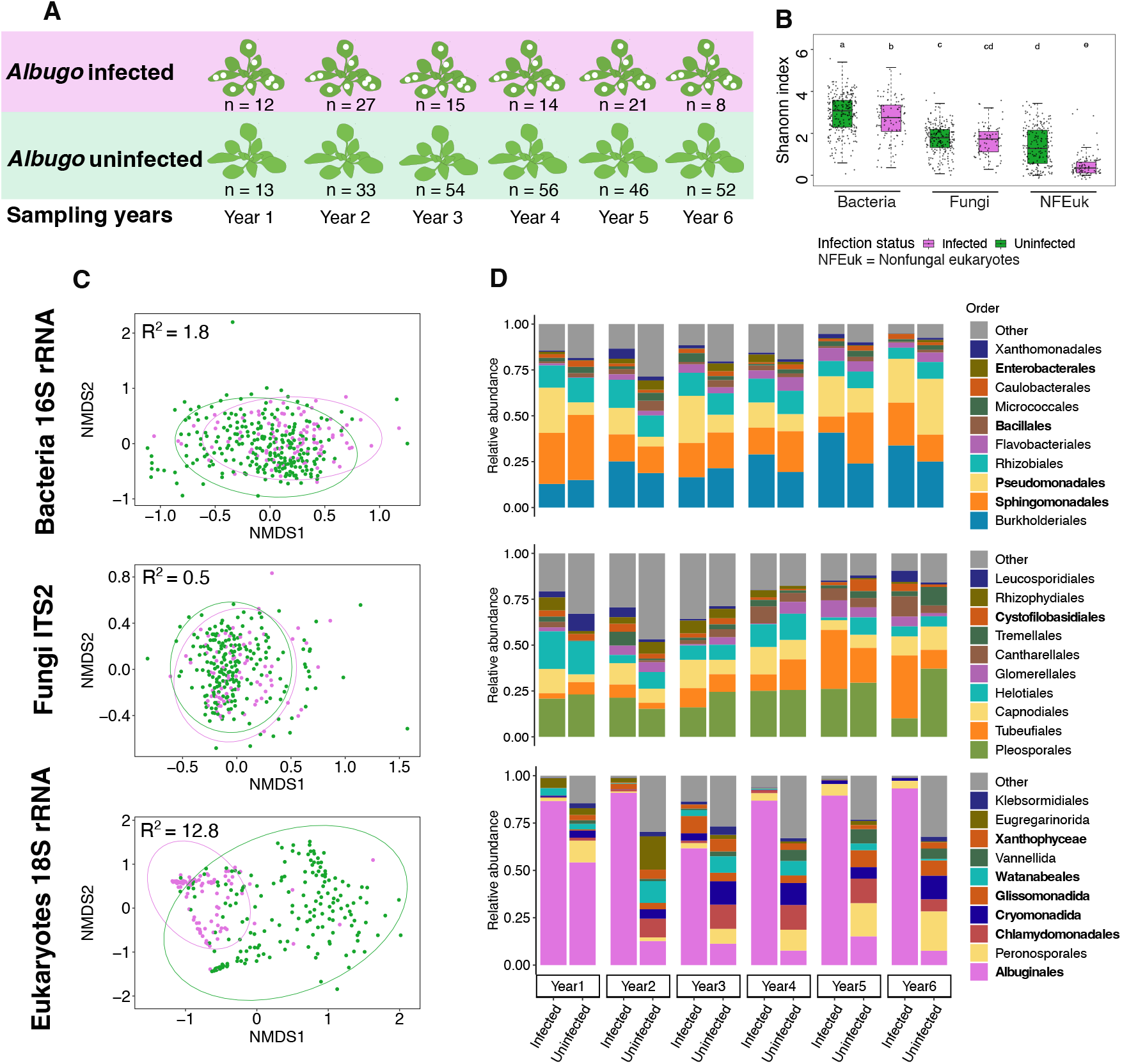
Diversity and composition of leaf microbial communities in *Albugo*-infected and uninfected plants. (A) The number of annually sampled plants per group (97 infected and 254 uninfected). (B) Alpha diversity is measured by Shannon’s H index and represents the within-sample diversity of infected and uninfected samples in bacteria, fungi, and nonfungal eukaryotic communities. Box plots display individual samples as dots. Different letters indicate statistically significant differences between groups (Tukey HSD’s test, *P <* 0.05). (C) Separation of infected (purple) and uninfected (green) samples using non-metric multidimensional scaling analysis (NMDS) based on Bray-Curtis dissimilarities. Each dot represents a single sample. (D) Histograms show the relative abundance of bacteria, fungi, and nonfungal eukaryotic communities at the order level, categorized by plant infection status (infected/uninfected) over six years. Taxa with significant differences between infection stages are indicated in bold (see Fig. S1).

A diversity analysis was conducted to compare the leaf-associated microbial communities between infected and uninfected plants. Alpha diversity (within-sample diversity, measured by Shannon’s index) demonstrated that, on average, infected plants exhibited a 1.1-fold and 2.8-fold reduction in bacterial and NFEuk community diversity, respectively, in comparison to uninfected plants (Tukey’s HSD test, *P <* 0.05). However, no significant differences were observed in fungal communities (Fig. 1B). A permutational multivariate analysis of variance (PERMANOVA) demonstrated that the ‘infection status’ of the plants explained 1.8% of the variation in bacteria, 0.5% in fungi, and 12.8% in nonfungal eukaryotes. These variations were further visualized using non-metric multidimensional scaling (NMDS), which revealed that the infected plants were most clearly separated from the uninfected groups in the NFEuk communities, followed by bacteria and less so in the fungal communities (Fig. 1C).

Differences between infected and uninfected communities were found to be associated with the enrichment of major microbial orders over sampling years (Wilcoxon test, *P <* 0.05) (Fig. 1D, see also Fig. S1). Among bacteria, *Sphingomonadales*, which has been demonstrated to be beneficial for plant health and productivity, [18] was 1.3 times more abundant in uninfected plants, whereas *Pseudomonadales* was 1.5 times more abundant in infected samples, based on a comparison of mean relative abundances. Among the fungal orders, only the Basidiomycete yeast *Cystofilobasidiales* exhibited a slightly higher significant abundance in infected plants (1.02 times more). Among nonfungal eukaryotic orders, as expected, the order *Albuginales* (including *Albugo*) showed a 6.7 times increase in infected plants. Interestingly, the most abundant orders of green algae (*Watanabeales, Xanthophyceae* and *Chlamydomonadales*) were 5.9-14.5 times more abundance in uninfected plants, similar to *Cercozoa* (*Glissomonadida* and *Cryomonadida*), with 3.0 and 7.6 times higher abundance in uninfected plants, respectively.

### Genotype variation in infected and uninfected *A. thaliana* correlates with microbiome diversity

To investigate whether leaf microbiome infection varies by host genotype, we used the whole genome sequencing data that was conducted to identify genotype clusters based on single nucleotide polymorphisms (SNPs) analysis [17]. Among the five identified clusters, three (clusters 1, 2, and 4) contain samples susceptible to *Albugo* infection. Clusters (2 and 4) exhibited significantly lower alpha diversity in NFEuk communities of infected plants as compared to uninfected clusters (Dunn test, *P <* 0.05) (Fig. 2A). A similar pattern was observed in the variability within clusters (i.e., how far each sample was from the group’s central point or centroid). Specifically, the clusters that were susceptible to infection showed less variability in this distance, meaning the samples in these groups were more similar to each other (Dunn test, *P <* 0.05) (Fig. 2B). However, no significant differences were found in fungal communities. Microbiome compositional variation, visualized using NMDS, showed that samples from cluster 4 more clearly separated the infected plants, while cluster 5 distinguished uninfected samples in the NFEuk community (Fig. 2C, PERMANOVA, *P <* 0.05). These patterns were less pronounced in bacterial communities (explaining 10.9% of the variation vs. 22.2% variation in NFEuk community) and were not significant in fungal communities (Fig. 2C).

**Figure 2.**
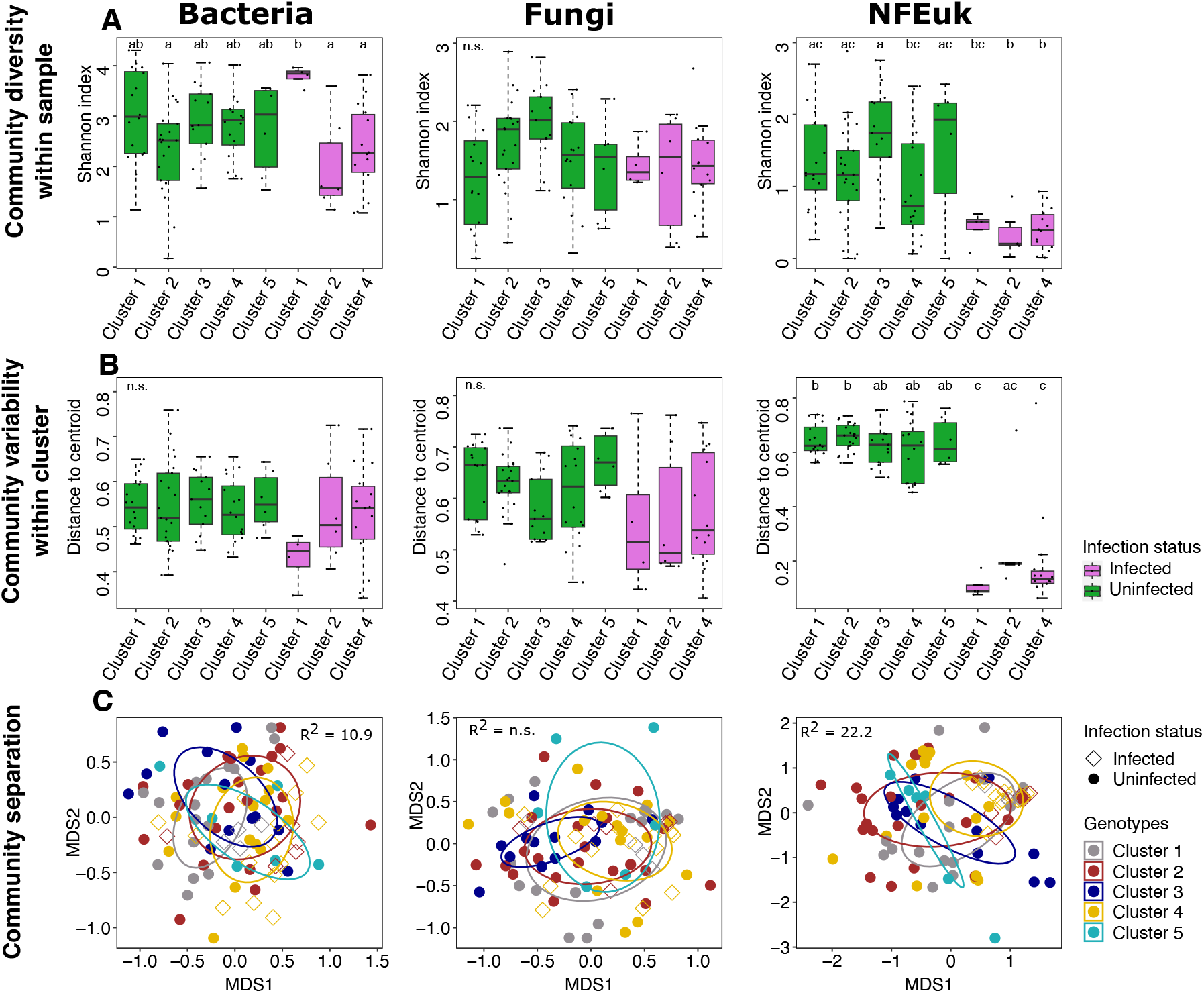
Diversity and separation of leaf microbial communities of infected and uninfected plants among plant genotypes. (A) Alpha diversity, measured by Shannon’s H index, represents the within-sample diversity in microbial communities across infected and uninfected samples of different genotype clusters. (B) Within-cluster variability, quantified as the distance of individual samples from the centroid of their respective genotype cluster, illustrates the variation among microbial communities within each cluster. Different letters indicate statistically significant differences between groups (Dunn test, *P <* 0.05). (C) Non-metric multidimensional scaling (NMDS) plots, based on Bray-Curtis dissimilarities, display the separation between infected and uninfected samples across genotype clusters. Explained variance (R^2^ values) from PERMANOVA models (Bray-Curtis dissimilarities), illustrating the impact of genotype clusters and infection status on the structure of leaf microbial communities.

Interestingly, the susceptible clusters were distributed across different sampling sites, suggesting that, in addition to host genotypes, other abiotic factors might contribute to plant susceptibility to infection. Notably, genotype cluster 5, which contained only uninfected plants, was exclusively found at the ERG site (Table. S1). Patterns related to sampling sites were observed: two sites (K69 and PFN) had no infected plants, while four sites (EY, WH, JUG, and ERG) contained both infected and uninfected plants (Fig. S2). Site EY exhibited lower alpha diversity in bacterial communities of infected plants compared to uninfected plants in the same group, while in NFEuk communities, three sites (EY, WH, and JUG) showed lower alpha diversity in infected plants (Dunn test, *P <* 0.05) (Fig. S2A). Regarding community variability, only site ERG showed reduced variability in infected plants within bacterial communities. In contrast, in NFEuk communities, the variability of infected plants was consistently lower than that of uninfected plants across all sites (Dunn test, *P <* 0.05) (Fig. S2B). The combination of infection status and sampling site explained 8.4%, 7.8% and 21.2% of the variation in microbial communities for bacteria, fungi and NFEuk, respectively (Fig. S2C).

### Identification of a microbial signature for predicting infected and uninfected leaves using machine learning models

We hypothesized that the plant microbiome is composed of distinct health-associated microbial communities (HCom) in uninfected plants and disease-associated microbial communities (DCom) in infected plants. These distinct microbial communities can serve as robust indicators of infection, enabling accurate discrimination between infected and uninfected samples. To investigate this hypothesis, we used machine learning classification models, including random forest (RF), support vector machine (SVM), and logistic regression (LR), which are well-known for their interpretability and multilayer perceptron (MLP) (Fig. 3A). The training phase used 70% of the sample set, consisting of 169 uninfected and 66 infected samples with 2,543 operational taxonomic units (OTUs). The remaining 30% (73 uninfected and 29 infected samples) served as the test set to assess predictive performance. Four different evaluation metrics were employed, resulting in accuracies ranging from 75% to 86% (Fig. 3C). The SVM and LR models achieved the highest accuracy of 85% and 86% respectively, with an area under the curve (AUC) of 93% and 94%, outperforming the MLP and RF (Fig. 3B and 3C). We then analysed the predictive role of each microbe by calculating and comparing the feature importance of all the OTUs in the trained classification models (SVM, RF, and LR) (Fig. 3A). Comparison of the three models revealed that they shared 2,253 OTUs, indicating consistent microbial signatures associated with both groups (infected/uninfected) that contributed significantly to the classification process (Fig. S3A). Using recursive feature elimination with cross-validation, we identified the most crucial OTUs for classification, referring to them as HCom and DCom. The results showed that RF had the highest accuracy, achieving 91% accuracy (Fig. S3B) with 40 selected OTUs (Fig. 4A). The LR model reached 87% accuracy with 4 selected characteristics (Fig. 4B). In contrast, the SVM reached 86% accuracy with 53 selected OTUs (Fig. 4C). It is interesting to note that four OTUs were shared by all three models that originated from LR, and 13 OTUs were shared by SVM and RF (Fig. 4 and Fig. S3C).

**Figure 3.**
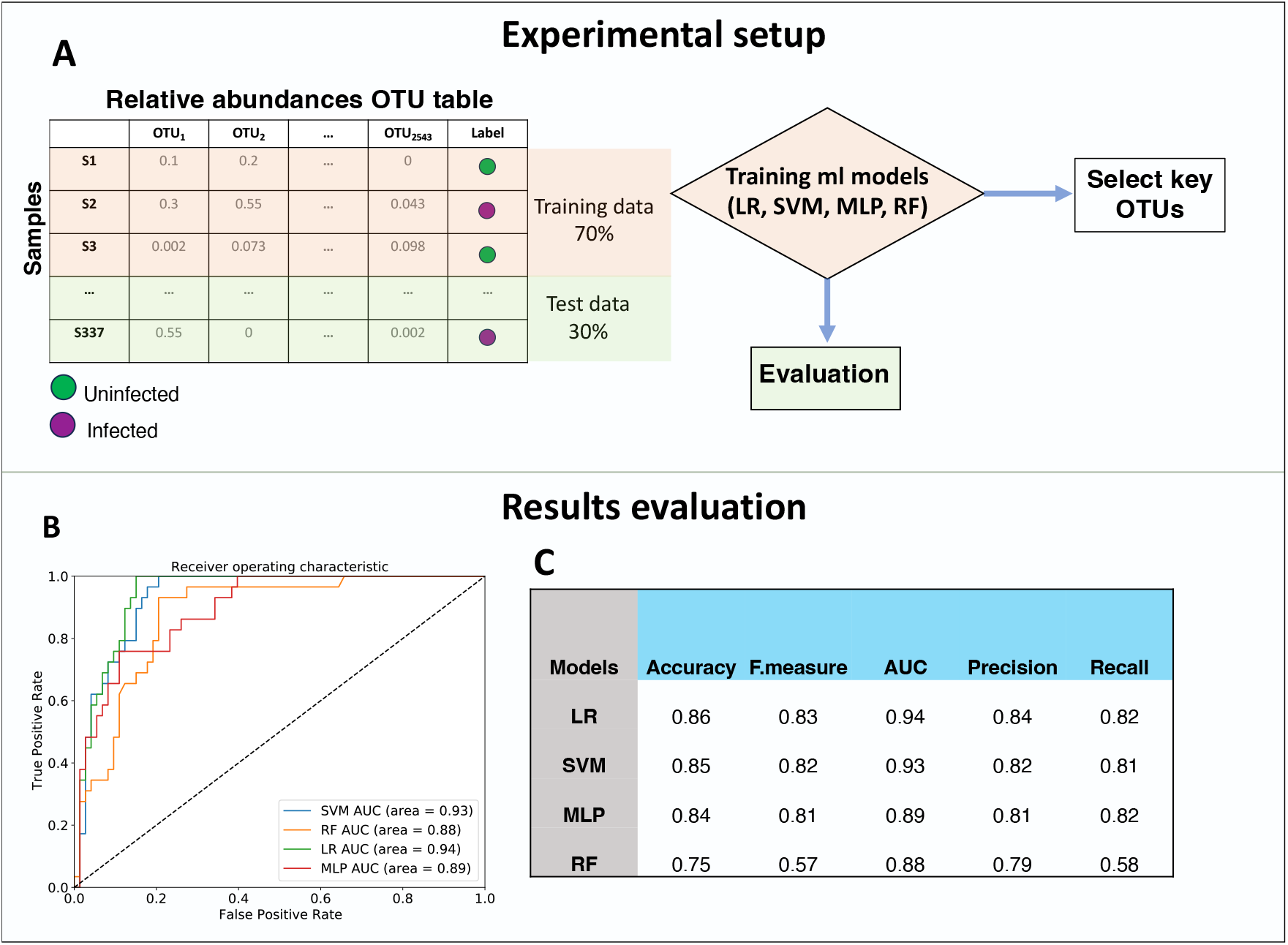
Classification of plant infection status and feature selection by machine learning classifiers. (A) Workflow illustrating the methodology employed to distinguish infected samples from uninfected ones using machine learning models. The objective was to classify leaves uninfected and infected based on the observed symptoms of *Albugo* infection, utilizing the relative abundance OTU table of bacteria, fungi, and nonfungal eukaryotes. Four machine learning classifiers, namely support vector machine (SVM), random forest (RF), logistic regression (LR), and multilayer perceptron (MLP), were trained on 70% of the samples. The trained models were evaluated using the remaining 30% of the dataset. Feature importance (to select key important OTUs for classification) was extracted from the trained models and using the recursive feature elimination method (Figure. 4). (B) Receiver operating characteristic (ROC) curves. The area under the curve (AUC) values indicate the ability of each classifier to distinguish between infected and uninfected samples, with higher AUC values indicating better performance. (C) Additional performance metrics (accuracy, f-measure, precision, and recall) for each classifier on the test set.

**Figure 4.**
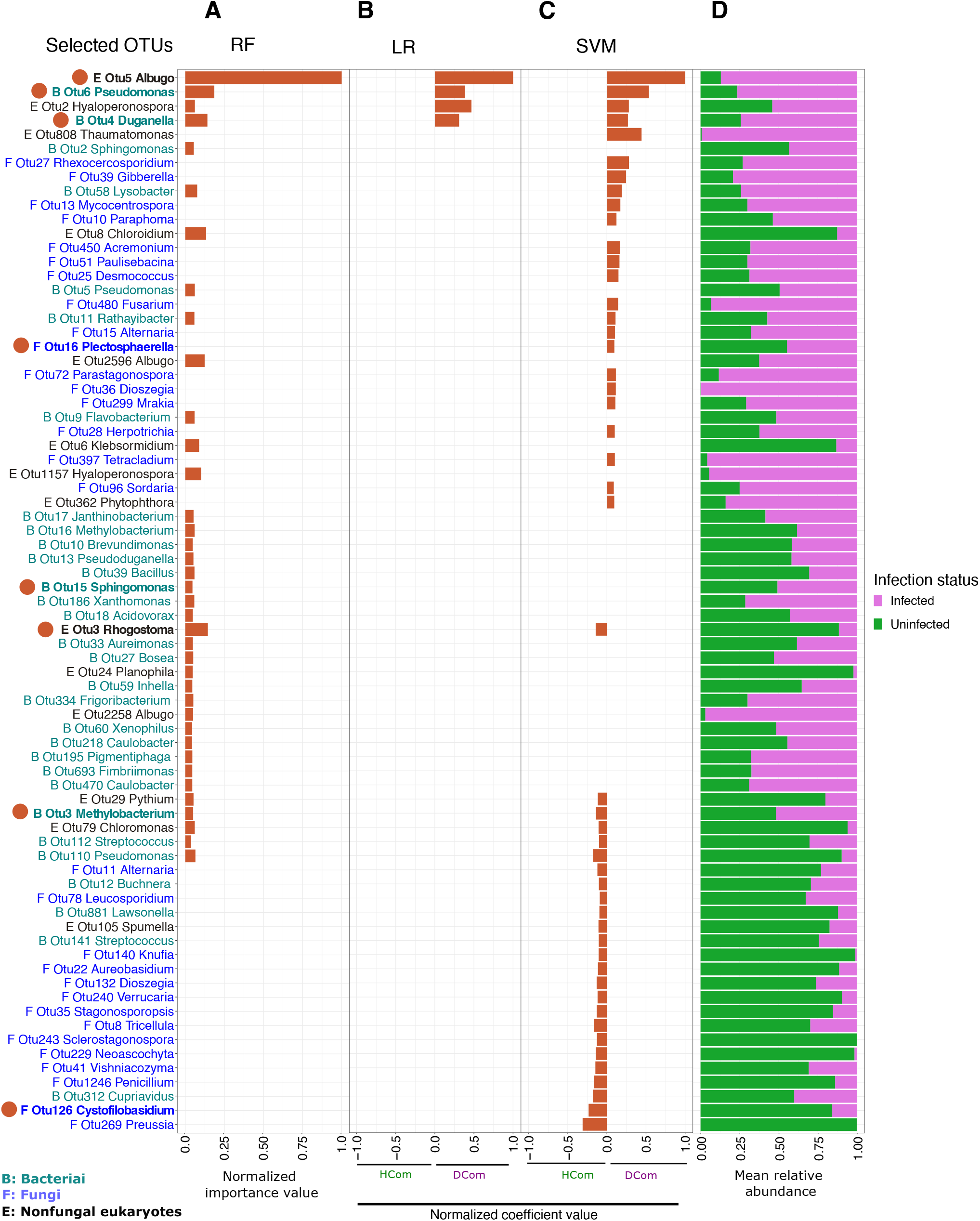
Selected HCom and DCom microbes as indicators of uninfected and infected samples. (A) Histogram showing the 40 OTUs selected by recursive feature elimination with k-fold cross-validation using random forest (RF). The x-axis indicates the importance of each OTU for classification (values are normalized between 0 and 1). (B) The 4 OTUs selected by recursive feature elimination with k-fold crossvalidation using logistic regression (LR). (C) Normalized coefficient values of the 53 OTUs selected by recursive feature elimination with k-fold cross-validation using support vector machine (SVM). Negative values (B and C) indicate OTUs with high scores in discriminating uninfected leaves (HCom), while positive values indicate high scores in discriminating infected samples (DCom). (D) Bar plots illustrating the aggregated relative abundances of OTUs in infected and uninfected samples. The microbes highlighted in bold (left) were selected for further experimental analysis.

Additionally, we hypothesized that HCom microbes can reduce the pathogenicity of *Albugo*, in contrast to DCom microbes. To test this hypothesis, we selected four candidate microbes from bacterial, fungal, and non-fungal eukaryotic groups within each category (HCom and DCom) (Fig. 4, microbes in bold). The SVM and LR models provide both positive and negative coefficients to determine microbial importance in classifying infected versus uninfected leaves. We assigned scores based on these coefficients, with negative scores indicating HCom OTUs as indicators of the uninfected class and positive scores representing DCom OTUs as indicators of the infected class (see Fig. 4B and 4C). However, the RF model only provides positive scores, necessitating further examination of the relationship between the selected OTUs and their respective classes (Fig. 4A). Other selection criteria included diverse representations of bacteria and eukaryotes, as well as laboratory availability. Among the HCom bacterial candidates, *Methylobacterium* OTU3 (*Methylobacterium goesingense*) and *Sphingomonas* OTU15 (*Sphingomonas melonis*) were selected, while *Cystofilobasidium* OTU126 (*Cystofilobasidium macerans*) and *Rhogostoma* OTU3 (*Rhogostoma epiphylla*) were selected for fungi and nonfungal eukaryotes, respectively. The Dcom candidates included *Duganella* OTU4 (*Duganella zoogloeoides*) and *Pseudomonas* OTU6 (*Pseudomonas viridiflava*) from the bacterial group, *Plectosphaerella* OTU16 (*Plectosphaerella niemeijerarum*) as fungal representative, and *Albugo* OTU5 (*Albugo*), representing nonfungal eukaryotes (Fig. 4A-C).

### Infection reduces microbial network complexity and increases compartmentalization in community structure

Microbial networks are valuable for identifying potential interactions among microorganisms within a given community. This is achieved by correlating the abundances of different species. Therefore, we investigated the microbial networks in both infected and uninfected samples. The network resulting from the uninfected samples showed 1.9 times greater number of nodes (OTUs) and 3.3 times more edges (connections between OTUs) when compared to the one generated from the infected samples (2,024 nodes and 73,511 edges vs. 1058 nodes and 22,089 edges, respectively) (Fig. 5A vs. Fig. 5B). Notably, the topological characteristics of the uninfected network showed higher degree (more connections between microbes) and closeness centrality (microbes more closely connected to others) values (*P <* 0.001) (Fig. 5C and 5D). To assess the effect of infection on the microbial communities’ compartmentalization, we compared the modularity of the constructed networks, a feature representing the degree of functional division and ecological niches within the microbial community [19]. The results showed that both networks contained a comparable number of modules, with the uninfected network having ten modules and the infected network having nine modules. However, the modularity value of the network derived from infected samples (0.31) was slightly higher than that of the network derived from uninfected samples (0.24) (Fig. 5B and 5A), which indicates that infected generated network is more segmented, suggesting a stronger tendency to division of microbial communities into distinct functional groups or ecological niches. Notably, all the modules contained OTUs from different taxonomic categories of bacteria, fungi, and nonfungal eukaryotes (Fig. 5E and 5F). We examined the microbial interactions of HCom and DCom OTUs in both uninfected and infected networks. Results showed that these OTUs displayed distinct connectivity patterns. In the network generated from uninfected samples, the OTUs formed 72 OTUs with 493 edges (Fig. S4A). In contrast, these OTUs exhibited fewer connections in the infected network, resulting in 64 nodes with 193 edges (Fig. S4B). Moreover, those OTUs are more sparsely distributed across different modules in the infected generated network compared to the uninfected network (Fig. S4D vs. Fig. S4C). The changes in connectivity patterns, particularly among the OTUs in the infected network, further highlight the reduced complexity and increased structural division in networks generated from infected samples.

**Figure 5.**
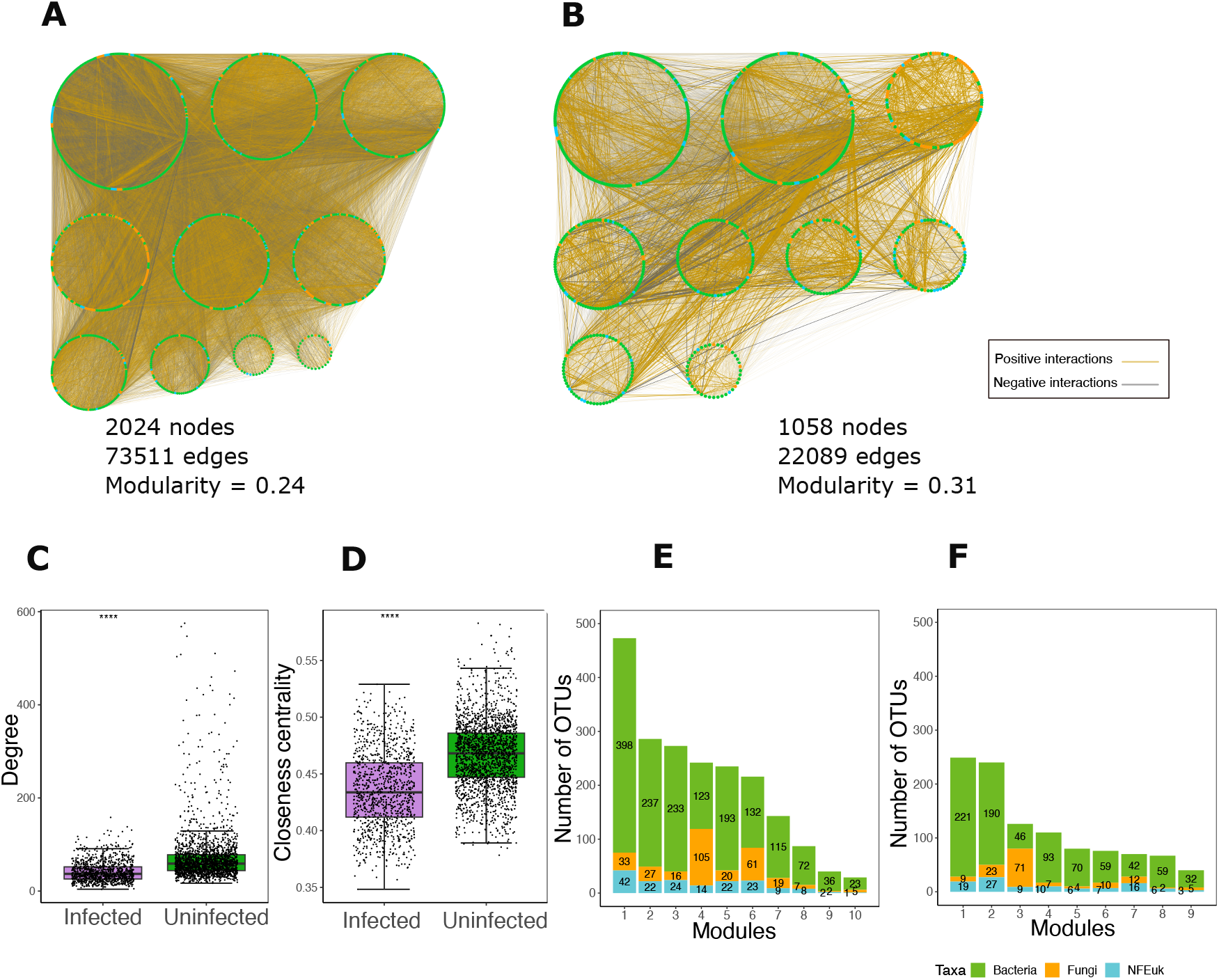
Changes in microbial co-abundance networks of infected and uninfected plants. Co-abundance networks for both uninfected (A) and infected (B) samples, where nodes (circles) represent OTUs and edges (the connection between OTUs) indicate correlations between these OTUs. Nodes are color-coded by microbial taxa and grouped based on modularity clustering. Box plots show features per node, i.e., degree (C) and closeness centralities (D) in infected and uninfected networks. Significance values indicate differences between groups based on the Wilcoxon test (\*\*\*\**P* ≤ 0.0001). Histograms illustrate the distribution of OTUs within modules for the network of uninfected (E) and infected (F) samples, respectively. These histograms are further color-coded to distinguish microbial taxa, with green representing bacteria, orange representing fungi, and blue representing NFEuk.

### HCom confer protection against *Albugo* infection to varying degrees

To investigate the protective effects of the selected microbes (Fig. 4) against the infection caused by *Albugo*, first, a mixture of *Albugo* and each of the four microbes from HCom was sprayed onto *Arabidopsis* leaves (Fig. 6A). The level of protection was determined by measuring the percentage of infected leaves (Fig. 6B). All four candidates significantly decreased the infection caused by *Albugo* (Dunn test, *P <* 0.05). *Cystofilobasidium* exhibited the most pronounced effect, reducing *Albugo* levels by an average of 73%. *Sphingomonas* caused a 66% reduction, followed by *Rhogostoma* and *Methylobacterium*, which resulted in 53% and 40% decreases in *Albugo* infection, respectively (Fig. 6B). These observations were further confirmed by quantitative polymerase chain reaction (qPCR) analysis, which demonstrated that samples exposed to the uninfected-associated microbes exhibited substantially lower amounts of *Albugo* as compared to control samples (Dunn test, *P <* 0.05), with the average biomass of *Albugo* ranging from 72% to 90% (Fig. 6C). These results demonstrate that all the selected candidates associated with uninfected plants, namely, *Cystofilobasidium, Methylobacterium, Rhogostoma*, and *Sphingomonas*, significantly decreased the infection levels of *Albugo*. The *Methylobacterium*-treated plants exhibited the highest plant biomass, with an average of 0.96 (g), compared to other treatments(Tukey’s HSD test, *P <* 0.05) (Fig. S5). Microscopy analysis revealed that *Rhogostoma* attached to the *Albugo* spores and feeds on free-living microbes in the environment (Fig. S6 and supplementary videos 1 and 2). These findings highlight the potential of HCom microbes in protecting against *Albugo* infection, with varying levels of effectiveness across different microbial taxa.

**Figure 6.**
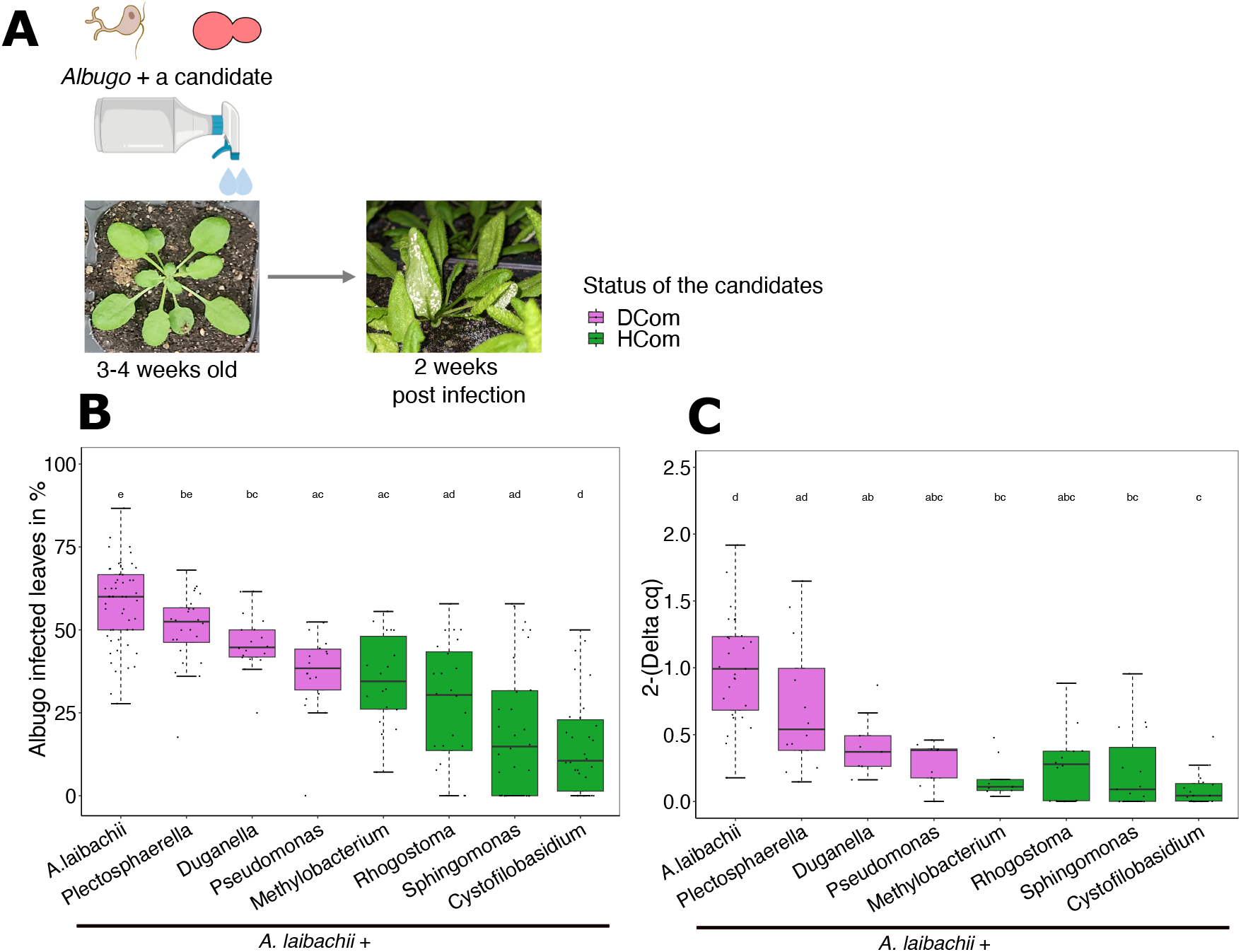
Effects of HCom and DCom on infection caused by *Albugo*. (A) Three to four weeks old *Arabidopsis* plants were co-inoculated with *Albugo* and each of the indicator microbes (HCom and DCom microbes) as identified in Figure 4. Symptoms were recorded 2 weeks after infection. (B) Box plots showing the percentage of leaves infected with *Albugo* in the presence of HCom (green) and DCom strains (purple). (C) Relative quantification of *Albugo* biomass in response to each indicator microbe was conducted through qPCR targeting the *Albugo* EF1-*α* gene and normalizing to the *A. thaliana* EF1-*α* gene. The relative biomass was then calculated via the ddcq method. Statistically significant differences between the groups were evaluated using the Dunn test, with different letters indicating significant differences (*P <* 0.05).

We then investigated the effect of DCom microbes on the pathogenicity of the *Albugo*. The *in-planta* infection assay demonstrated that *Plectosphaerella* had no significant effect on the infection level of *Albugo* (Dunn test, *P >* 0.05). However, *Pseudomonas* and *Duganella* caused a decrease in infection of 36% and 20%, respectively (Dunn test, *P <* 0.05) (Fig. 6B). Likewise, qPCR outcomes supported the observed phenotype: the biomass of *Albugo* in the control group exhibited no significant changes in comparison with *Plectosphaerella* (Dunn test, *P <* 0.05), whereas *Pseudomonas* and *Duganella* caused a 72% and 58% reduction in *Albugo* biomass, respectively (Fig. 6C). The findings reveal that while some DCom microbes are effective in reducing *Albugo* infection levels, their protective effects are less significant compared to those of HCom microbes.

To evaluate the specific effects of each microbe on plant health independently, we performed spray experiments on gnotobiotic plants, isolating the impact of individual microbes without the influence of other microbial interactions (Fig. S7). Three distinct phenotypes were observed. Plants colonized with the DCom bacteria *Duganella* and *Pseudomonas* exhibited high mortality rates, with 10.0% and 11.3% survival, respectively, within three weeks post-colonization. As expected, the filamentous pathogenic *Plectospherella* and *Albugo* caused characteristic infection symptoms, namely, brownish leaves and roots and white rust disease, respectively. Forty-nine percent of the plants treated with HCom *Cystophilobasidium* survived three weeks postcolonization. The healthiest plants were those colonized by the remaining HCom microbes—*Sphingomonas, Methylobacterium*, and *Rhogostoma*-with over 88% of the plants showing no discernible negative consequences. Overall, these results indicate that most HCom microbes lead to healthier plants than DCom microbes, underscoring the importance of specific microbial candidates in enhancing plant vitality and resistance to pathogen *Albugo*.

## Discussion

In this study, we identify microbial signatures that distinguish between infected and uninfected plants and explore their potential for developing effective probiotics to promote plant health. The management of plant health through natural probiotics has gained significant ecological and economic interest. Various microbes and synthetic microbial communities have been found to increase plant resistance to pathogens under laboratory conditions [12, 20]. Pathogens are known to impact the phyllosphere microbiome to establish their niche [21]. Recent studies show that pathogens, including *Verticillium* and *Albugo*, release effector proteins to manipulate the microbial landscape, affecting microbiome composition and function [16, 22, 23]. However, there is a gap in understanding these interactions under natural field conditions, where pathogens face a complex and heterogeneous host microbiome and abiotic stressors [17, 24]. Our study addresses this by investigating microbial communities associated with *A. thaliana* under natural field conditions over six years, focusing on changes in the presence or absence of *Albugo* infection.

Agler et al. [15] reported that the diversity of bacterial communities was lower in *Albugo*-infected plants. Extending this observation, our analysis of six-year time series data, along with 18S eukaryotic data, revealed that not only bacterial and fungal diversity affected by infection, but there was an even more pronounced loss of diversity within 18S nonfungal eukaryotic groups (Fig. 1). This loss was observed in 3 out of 5 host genotypes that showed susceptibility to infection, leading to up to a 22% variation in the microbiome composition (Fig. 2). We attribute this reduction in diversity to a significant increase in the pathogen population, which can disrupt the balance of the established microbial community [10, 25]. In the case of *Albugo* infection, this imbalance may result from the pathogen’s efficient suppression of host defenses, allowing nonhost pathogens to proliferate [26], or from the release of microbial-modulating effector proteins [16]. Both mechanisms likely contribute to the establishment and maintenance of the pathogen’s niche within the host.

To identify signatures, the pathogen *Albugo* imposes on the *A. thaliana* microbiome, we used machine learning prediction models to identify non-linear relationships and manage the complexity of the high-dimensional data [27]. Therefore, we conducted a systematic analysis of variations between uninfected and infected plant statuses, achieving highly accurate classifications up to 91% (Fig. 3 and Fig. S3). These findings suggest the presence of predictive microbial signatures in these groups. Using the feature selection technique, we pinpointed 3.1% of OTUs, including bacteria, fungi, and nonfungal eukaryotes, as key discriminators between infected and uninfected plants, corresponding to HCom and DCom microbes (Fig. 4 and Fig. S3). These results highlight that, despite the vast microbial diversity, only a small subset is significantly associated with plant health outcomes. Interestingly, we found that HCom and DCom microbes are distributed across various network modules (Fig. S4). Modularity in microbial interaction networks can indicate diverse habitats, varying selective pressures, and phylogenetic clustering of related species [28], highlighting the crucial role of these microbes in the functionality of different community modules within the overall microbial community.

Building on these findings, we aimed to test whether microbial isolates matching the taxa identified by our machine learning prediction models could promote health or disease status in field conditions. Here, we could show that the predicted HCom and DCom microbiome comprises distinct phylogenetic groups of bacteria, fungi, and nonfungal eukaryotes (Fig. 4). Interestingly, some taxa were represented in both infected and uninfected samples. These included known pathogens like the fungal Ascomycetes genus *Alternaria*, saprophytic fungi such as the Basidiomycete yeast genera *Dioszegia* and *Cystofilobasidium* and the Ascomycetes order *Heliotales*. In addition, the bacterial genera *Pseudomonas* and *Sphingomonas* were also present in both groups. *Pseudomonas* is particularly notable for its ambivalent behavior, with a broad range of sublineages that can be either pathogenic or protective [29], either through microbe-microbe interactions or host interactions ([30]. This diversity was reflected in our dataset, where *Pseudomonas* OTU5 and OTU6 were predicted to be associated with infected samples, while *Pseudomonas* OTU110 was associated with health. A similar diversity has been reported for *Sphingomonas* [31], which, similar to *Pseudomonas*, can protect *A. thaliana* from pathogenic bacteria [32].

Given their relevance to *A. thaliana* health, we selected *Pseudomonas* and *Sphingomonas* isolates for testing. Our tests confirmed that the *Pseudomonas* isolate was pathogenic, while the *Sphingomonas* isolate did not harm the plant when colonizing the host alone (Fig. S7). From our HCom microbes, *Methylobacterium* and *Sphingomonas* showed further robust suppressive effects (Fig. 6). This is consistent with previous observations that *Sphingomonas* is able to suppress *Pseudomonas syringae* disease symptoms [32]. Further to the genera of *Spnignomonas* and *Pseudomonas*, members of the *Methylobacterium* genus and the family *Oxalobacteriaceae* are highly adapted to live on plants with high abundance and diversity [33, 34]. While *Methylobacterium* was predicted to be HCom in our analyses, the *Oxalobacteriaceae* genus *Duganella* was strongly associated with *Albugo* infection. Our experiment validated that the *Duganella* isolate exhibited pathogenic behavior (Fig. S7).

Among the HCom selected microbes, the basidiomycete yeast *Cystofilobasidium* exhibited the strongest protective effects (Fig. 6). *Cystofilobasidium* is particularly noteworthy, as several basidiomycete yeasts have been empirically identified as biocontrol agents against postharvest diseases in various fruits. This includes our computationally identified yeasts *Vishniacozyma* [35] and *Leucosporidium* [36], as well as our experimentally validated yeast, *Cystofilobasidium* [37]. Beyond yeast, another crucial group influencing plant health is the *Cercozoa*, which plays a significant role in shaping bacterial and fungal communities through selective predation [38, 39]. Our study demonstrates that the HCom *Rhogostoma* can suppress infection. As primary microbial predators, *Rhogostoma* directly targets bacterial and fungal pathogens, exerting a consumptive effect on various pathogenic strains [40, 41]. The reduction in infection levels may occur through direct consumption of *Albugo* zoospores or by preying on bacterial and fungal species that facilitate *Albugo* infection. Additionally, *Rhogostoma* may indirectly contribute to plant health by promoting beneficial microbes and enhancing interactive activities, such as biofilm formation, as has been recently shown for the protist *Cercomonas lenta* in the rhizosphere [42].

In summary, our findings support the concept of functional similarity [43], where diverse microbial taxa, including bacteria, fungi, and nonfungal eukaryotes, share overlapping ecological roles that contribute to plant health. This concept suggests that while different microbial species can perform similar functions, their effectiveness may vary depending on environmental conditions. Therefore, if abiotic factors (e.g., radiation and humidity) become unfavorable for one group, others can compensate, ensuring continued protection against pathogens [17]. Leveraging functional similarity presents promising opportunities for biocontrol methods to reduce pathogen pressure across diverse environmental conditions. As climate change increases the frequency of environmental fluctuations, leading to greater stress on plants, enhancing environmental resilience through the use of robust probiotics will be essential for future food security.

## Method

### Diversity analysis

The OTU tables for bacteria, fungi, and nonfungal eukaryotes, along with the corresponding genotype clusters for each sample, were obtained from the study by Mahmoudi et al. [17]. The OTU tables were modified by excluding epiphytic samples and those with fewer than 50 reads. Subsequently, OTU abundance tables were utilized to compute Shannon’s H diversity index using the ‘estimate-richness’ function in the Phyloseq R package [44] for estimating alpha-diversity. To assess between-sample diversity, relative abundance OTU tables were computed and transformed using log10 (x + 1) before calculating Bray-Curtis dissimilarities, which were then employed for nonmetric multidimensional scaling ordination (NMDS) using the ‘ordinate’ function in Phyloseq [44] R package. Three PERMANOVA analysis on Bray-Curtis dissimilarities were conducted to identify the primary factors (‘infection stages’, ‘combined infection stages and genotype’ and ‘combined infection stages and sampling sites’) influencing the leaf microbiome’s structure, utilizing the ‘adonis2’ function in the Vegan package [45] with 10,000 permutations (P *<* 0.05). A beta-dispersion analysis on Bray-Curtis dissimilarities was conducted to compare sample-to-sample variation within each group (genotype clusters and sampling sites) (multivariate homogeneity of group dispersion analysis, “betadisper”; Vegan package). The means were compared using the nonparametric multivariate test for multiple groups (‘dunnTest’ function in the FSA package [46], with BenjaminiHochberg adjusted P-values *<* 0.05), and the nonparametric ranked test for two groups (‘wilcox.test’ function in the stats package [47], P *<* 0.05). All analyses were conducted in R (version 4.1.2) [48].

### Machine learning analysis

The OTU tables of bacteria, fungi, and nonfungal eukaryotes were converted into relative abundance tables and merged into a single table. The OTU tables were further filtered to retain only those OTUs present in at least five samples (2543 OTUs and 337 samples). A processed dataset was created using two binary labels, ‘infected’ and ‘uninfected,’ corresponding to the phenotype of the collected plants. Plants were labeled ‘infected’ if white rust disease caused by *Albugo* was observed and ‘uninfected’ if it was not. The scikit-learn package in the Python programming environment was used to train and validate machine learning classifiers [49]. First, samples were divided into two parts: training (70%) and testing (30%) (train_test_split function, test_size=0.3, shuffle=True). Subsequently, machine learning models were trained. The trained models included Support Vector Machine (svm.SVC function, kernel=‘linear’), Random Forest (RandomForestClassifier function, n_estimators=1000, min_samples_split=2), multilayer perceptron (MLPClassifier function, solver=‘adam’, alpha=1e-5, random_state=1, learning_rate=“adaptive”, max_iter=500, hidden_layer_sizes=(100,100,100)), and Logistic Regression (LogisticRegression function, default parameters). The labels of the test sets were predicted (model.predict function). The prediction results were compared with the actual labels of the samples to calculate the models’ performance in terms of accuracy (accuracy_score function), f-measure (f1_score function, average=‘macro’), precision (precision_score, average=‘macro’), and recall (recall_score function). The area under the curve (AUC) was calculated using false positive and true positive rates (roc curve function) and plotted using the matplotlib package. The importance of each OTU in the trained models was obtained using model.coef_ (for Support Vector Machine and Logistic Regression) and model.feature_importances_ (for Random Forest). Recursive feature elimination was used to identify the most important OTUs (HCom and DCom) for classifying groups (RFECV function, step=1, cv=total number of samples, shuffle=True, scoring=‘accuracy’, min_features_to_select=1).

### Microbial network analysis and properties

To construct microbial correlation networks, samples of infected and uninfected plants were separated, and the OTU tables encompassing bacteria, fungi, and nonfungal eukaryotes were merged to conduct a comprehensive examination of microbial interactions. The OTU tables were then filtered to retain only OTUs present in at least five samples, resulting in 2,543 OTUs across 242 samples for uninfected plants and 1,058 OTUs across 95 samples for infected plants. These filtered OTU tables were used to calculate correlations using the SparCC algorithm[50], which uses Aitchison’s log-ratio analysis and is specifically designed to handle compositional data with high sparsity. The SparCC correlation scores were computed on the FastSpar platform [51] with default parameters. Pseudo P-values were generated through 1000 bootstraps to assess statistical significance. For further analysis, only correlations meeting the criteria of P*<*0.01 were included. Modularity analysis was performed using Python’s networkx package (version 3.1) [52]. Community detection was applied using the Louvain Community Detection Algorithm (community.community_louvain.best_partition function)) [53] the modularity score was calculated on detected modules (and community.community_louvain.modularity functions). We used Cytoscape (version 3.7.1) [54] to visualize and analyze the microbial interaction networks.

### Infections of *A. thaliana* leaves and quantification of *Albugo* biomass by qPCR

Overnight liquid cultures of bacteria and yeast were diluted in the fresh medium until they reached an OD_600_ of 0.2. The resulting cultures were then centrifuged at 1200 g for 5 minutes, and the resulting pellets were resuspended in MgCl_2_. A spore and cell concentration of 25 × 10^4^ spores/mL or cells/ml was prepared for *Plectospherella* and *Rhogostoma*. A spore solution of *Albugo* was prepared by collecting *Albugo laibachii* Nc14, which had previously been cultured on *A. thaliana* Ws-0. Water was added to the leaves, and the samples were kept on ice for 1 hour before filtering. The number of cells (*Rhogostoma*) or spores (*Plectospherella* and *Albugo*) was measured by taking 50 *μ*L of the solution, placing it on a hemocytometer, and examining it under an epifluorescence Axiophot microscope. Approximately 4-5 mL of each sample were carefully combined with 5-6 mL of *Albugo* solution (25 × 10^4^ spores/mL) and evenly applied to 4-5 weeks old *A. thaliana* seedlings (Ws-0 accessions) using airbrush guns. Following two weeks, leaf disease symptoms were assessed, differentiating between infected and uninfected leaves, and quantified as a percentage, and plants’ fresh weights were measured. The leaves were then stored at −80°C. DNA extraction was performed using FastDNATM Spin Kit for Soil (MP Bio) as described in the manufacturer’s protocol. For qPCR, a mixture was prepared consisting of 7.5 *μ*L of SYBR Green supermix, 5 *μ*L of DNA (approximately 50 ng), 1.9 *μ*L of NFW, and 0.3 *μ*L of forward and reverse primers (10 *μ*M each), resulting in a total reaction volume of 15 *μ*L. The sample measurements were triplicated using a Bio-Rad CFX Connect real-time PCR detection system. The quantification of *Albugo* DNA relative to the plant DNA was determined using the following oligonucleotide sequences: *A. thaliana* EF1-*α*: 5’- AAGGAGGCTGCTGAGATGAA-3’, 5’-TGGTGGTCTCGAACTTCCAG-3’; *Albugo* EF1-*α*: 5’-GTGTTCTGCACATCCACACC-3’, 5’-GACCTTGACGGATGAAAGGA- 3’. Cq values obtained during the amplification of oomycete DNA were subtracted from DNA amplicons of *A. thaliana* (called ddCq). Then, the relative biomass of *Albugo* for each group was determined using the formula 2^−ddCq^. These results were further normalized by comparing the *Albugo* biomass in each experiment (x) to the control. Specifically, we calculated the ratio of the ddCq for any treatment in experiment x to the average ddCq of the *Albugo* control in the same experiment (experiment x).

### Infections of *A. thaliana* leaves with individual strains in sterile conditions

Following the protocol of Eitzen et al., [23], sterilized *A. thaliana* seeds (Ws-0 accessions) were sown on 1/2 strength Murashige Skoog (MS) medium and incubated for 3-4 days in a cold room (4C, darkness). The MS plates were then placed in growth chambers set to 22°C with a short-day light cycle (8 hours of light) and 33-40% humidity. The seedlings were grown under these conditions for 4-5 weeks before inoculation. Overnight liquid cultures of bacteria and yeast were diluted in the fresh medium until they reached an OD_600_ of 0.2. The resulting cultures were then centrifuged at 1200 g for 5 minutes, and the resulting pellets were resuspended in MgCl_2_. A spore concentration of 25 × 10^4^ spores/mL was meticulously prepared for *Plectospherella* and *Rhogostoma*. Five hundred ul of each culture was evenly sprayed onto 4-5-week-old *A. thaliana* seedlings using airbrush guns. Phenotypes of plants were scored after 2-3 weeks post-infection.

### Microscopical observations

A mixture of *Albugo* spores and *Rhogostoma* cells, each at a concentration of approximately 100,000 cells/ml, was prepared and placed in a plastic petri dish. Observations were made under an epifluorescence Axiophot microscope (Zeiss, up to 64x magnification) and/or inverted microscope (Zeiss LSM880) 2 to 8 days post-mix.

## Availability of data and material

OTU tables and scripts are available here https://gitlab.plantmicrobe.de/maryam_mahmoudi/HealthMarkers

## Author Contributions

MM, KN, and EK devised the study. MM and LMT performed the experiments. MM, JA and YH interpreted the data. MM wrote the scripts and visualized the data. MM, JA, YH, KN and EK contributed to writing and preparing the manuscript. All authors read and approved the final manuscript.

## Acknowledgements

We thank Dr. Libera Lo Presti for the critical reading of the manuscript. We further thank the de.NBI Cloud Storage and bwForCluster BinAC Tübingen provide resources for storing and analyzing the data.

## Supplementary figures

**Supplementary figure 1.**
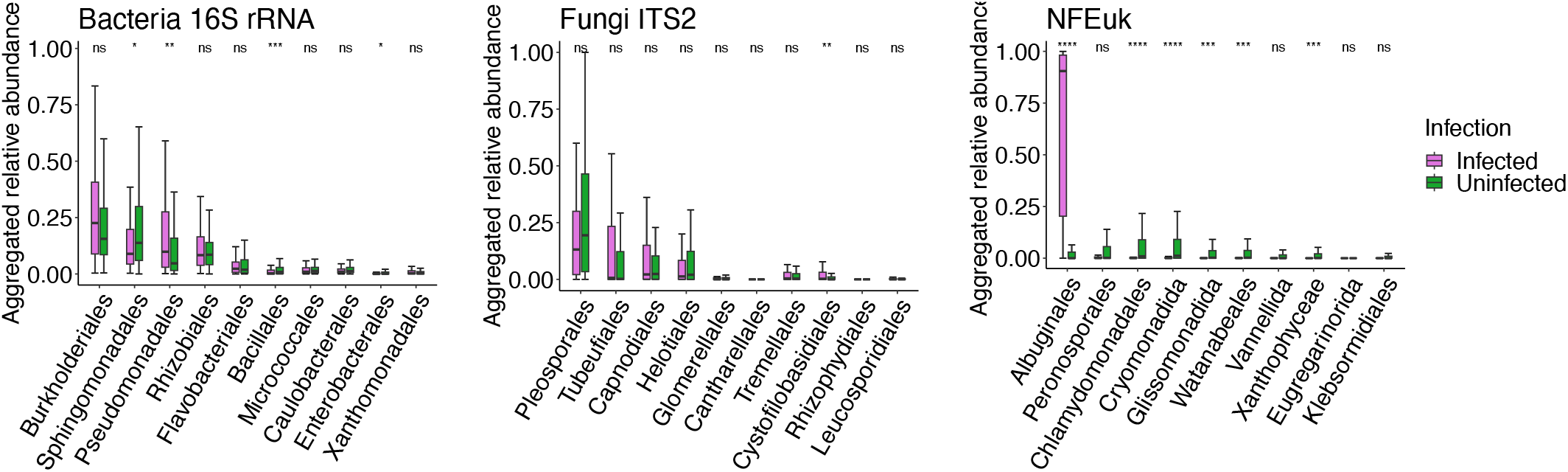
Changes in highly abundant microbial taxa colonizing *A. thaliana*’s in infected and uninfected leaves. Box plots (green = uninfected, purple = infected) show the relative abundance of the orders of bacteria, fungi and nonfungal eukaryotes in individual samples aggregated by ‘infection status’. Significance values between groups are based on Wilcoxon’s test: n.s. (*P >* 0.05), * (*P* ≤ 0.05), ** (*P* ≤ 0.01), *** (*P* ≤ 0.001), and **** (*P* ≤ 0.0001).

**Supplementary figure 2.**
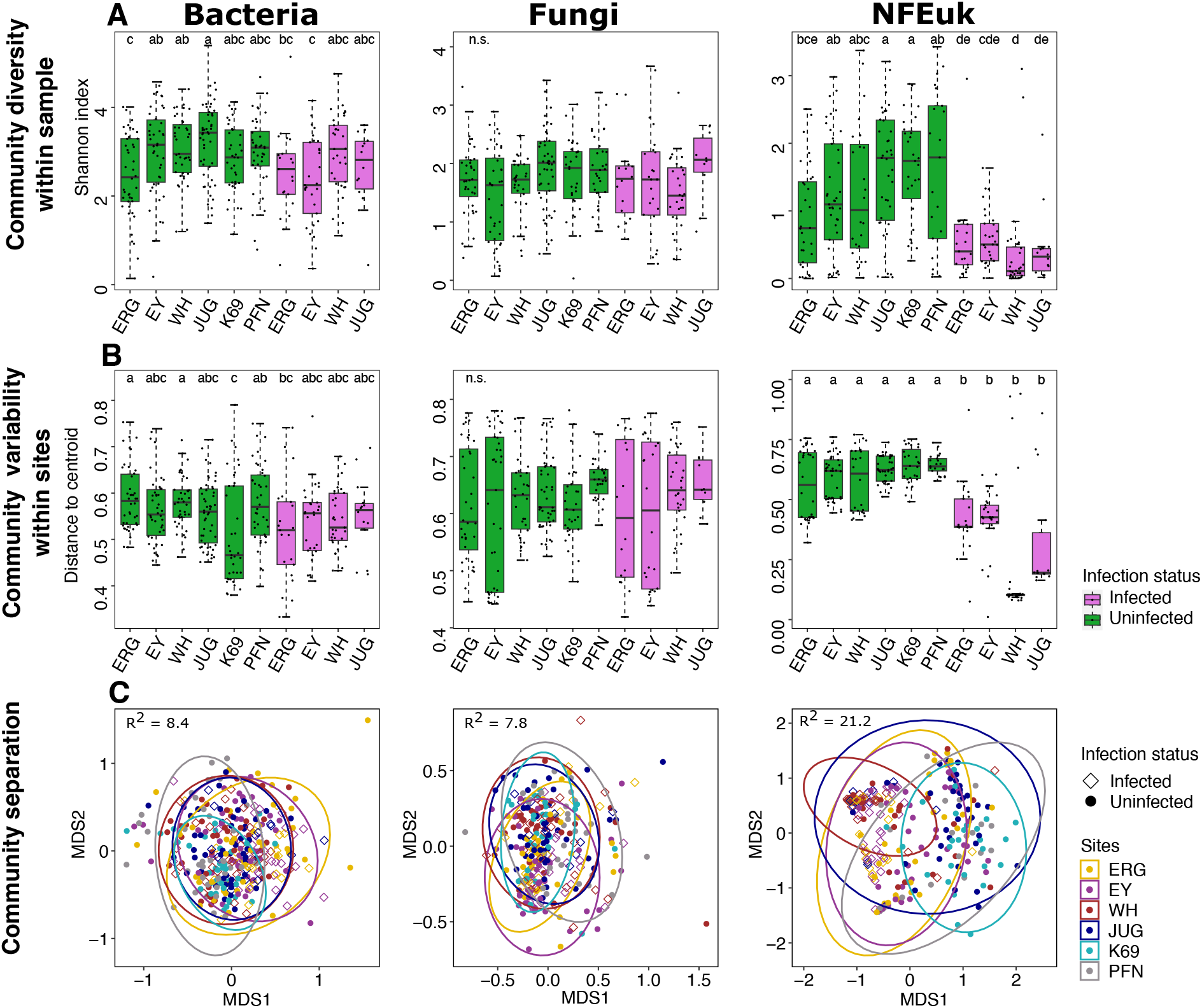
Diversity and variability of leaf microbial communities of infected and uninfected plants among sampling sites. (A) Alpha diversity, measured by Shannon’s H index and representing the within-sample diversity of infected and uninfected samples among sampling sites in microbial communities. (B) Within-site variability (distance to the group centroid). Different letters indicate statistically significant differences between groups (Dunn test, *P <* 0.05). (C) Nonmetric multidimensional scaling (NMDS) plots, based on Bray-Curtis dissimilarities, display the separation between infected and uninfected samples across sampling sites. Explained variance (R^2^ values) from PERMANOVA models (Bray-Curtis dissimilarities), illustrating the impact of sampling sites and infection status on the structure of leaf microbial communities.

**Supplementary Figure 3.**
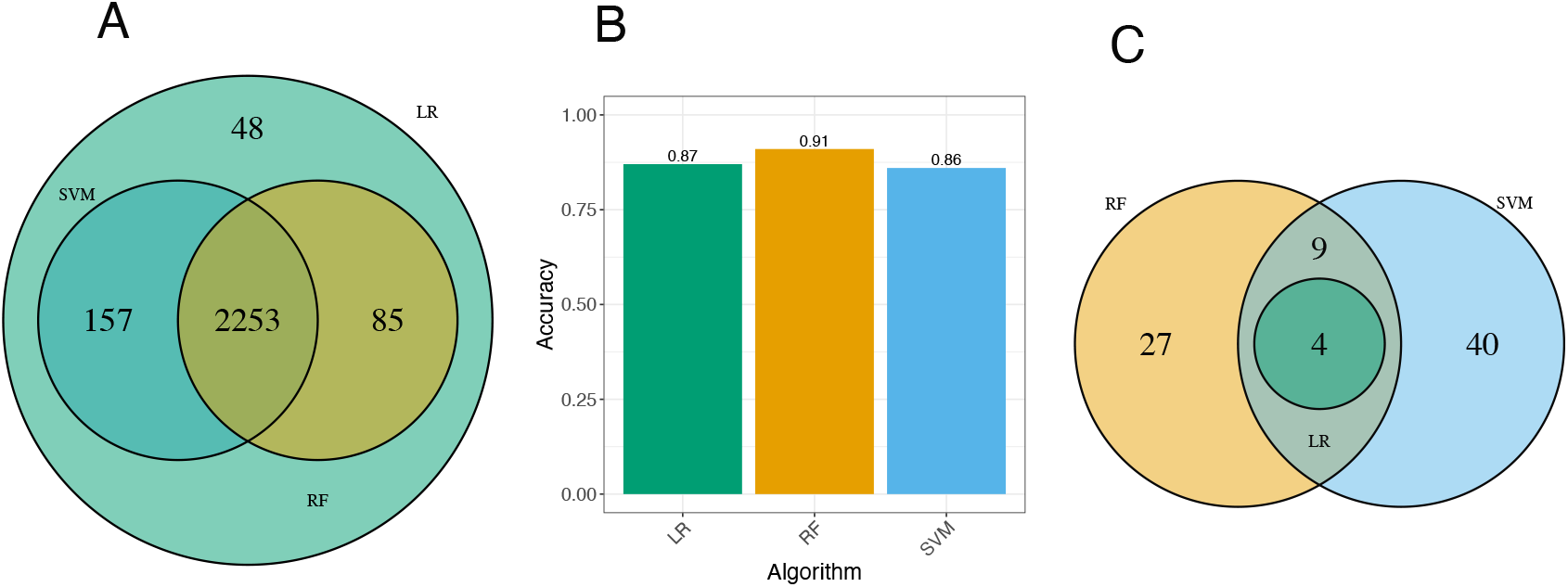
Comparing the importance of microbes for classifying leaves in uninfected and infected using machine learning models. To discriminate infected from uninfected samples based on microbial signatures, four classification models were trained (Figure. 3). (A) The Venn diagram represents the number of common microbes with absolute scores greater than 0 among different models during the training phase. (B) Bar plots represent the accuracy of models trained using recursive feature elimination. (C) The Venn diagram represents the number of shared microbes with absolute scores higher than 0 after performing recursive feature elimination.

**Supplementary Figure 4.**
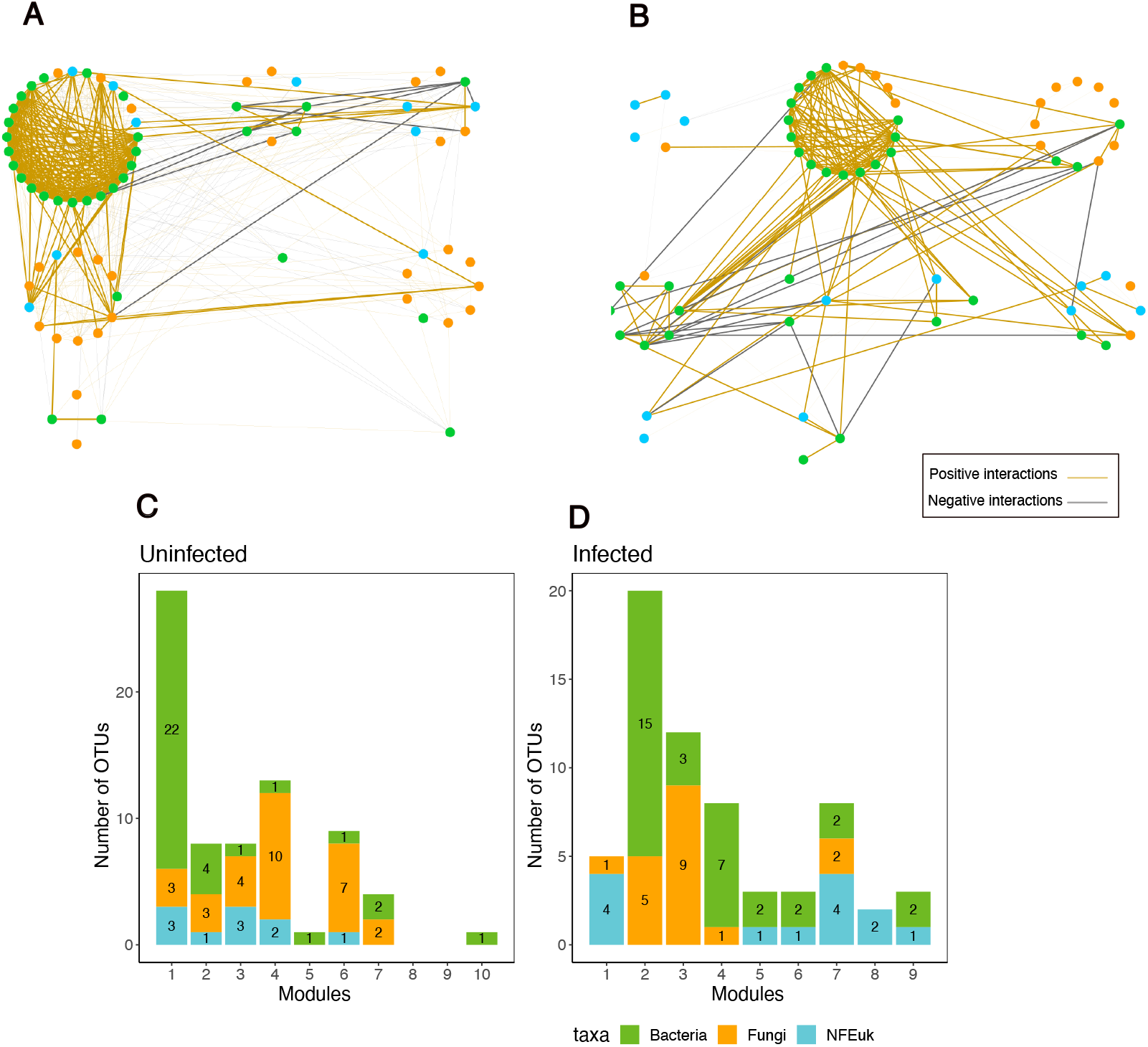
Changes in the co-abundance interactions of HCom and DCom OTUs. Interactions between HCom and DCom OTUs in the network constructed for uninfected (A) and infected (B) samples. OTUs are clustered by module and colored by taxa. Edges are colored according to the correlations: positive correlations are colored in orange, and negative correlations are colored in gray. Histograms (C and D) show the OTU distribution within modules for the networks of uninfected and infected samples, respectively. These histograms are further color-coded to distinguish microbial taxa: green represents bacteria, orange represents fungi, and blue represents nonfungal eukaryotes.

**Supplementary Figure 5.**
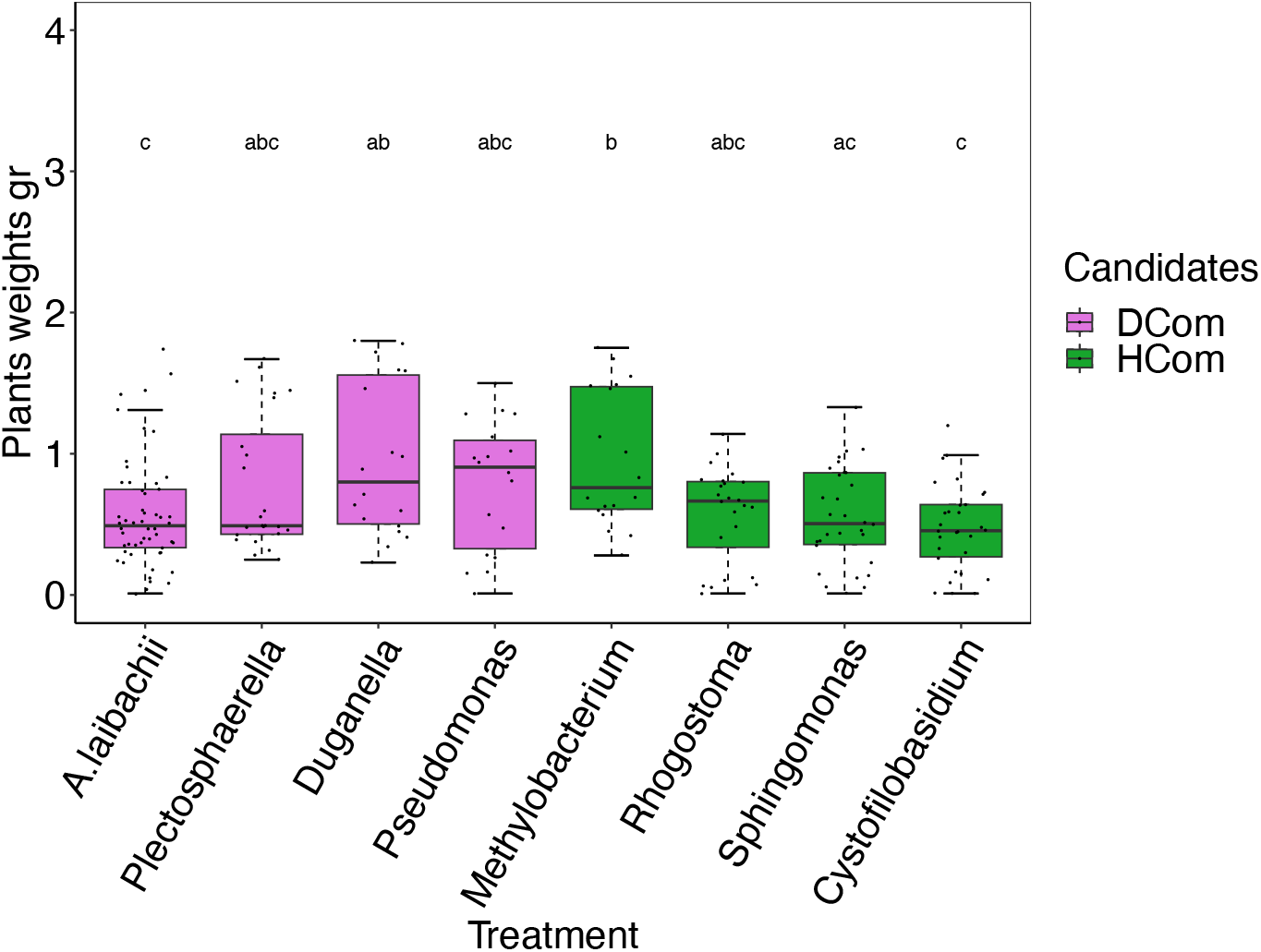
Fresh weight of plants inoculated with selected strains (see Figure. 6). Box plots showing the weights of leaves infected with *Albugo* in the presence of HCom strains (green) and DCom strains (purple). Statistically significant differences between the two groups were evaluated using Tukey’s HSD test, with different letters indicating significant differences (*P <* 0.05).

**Supplementary Figure 6.**
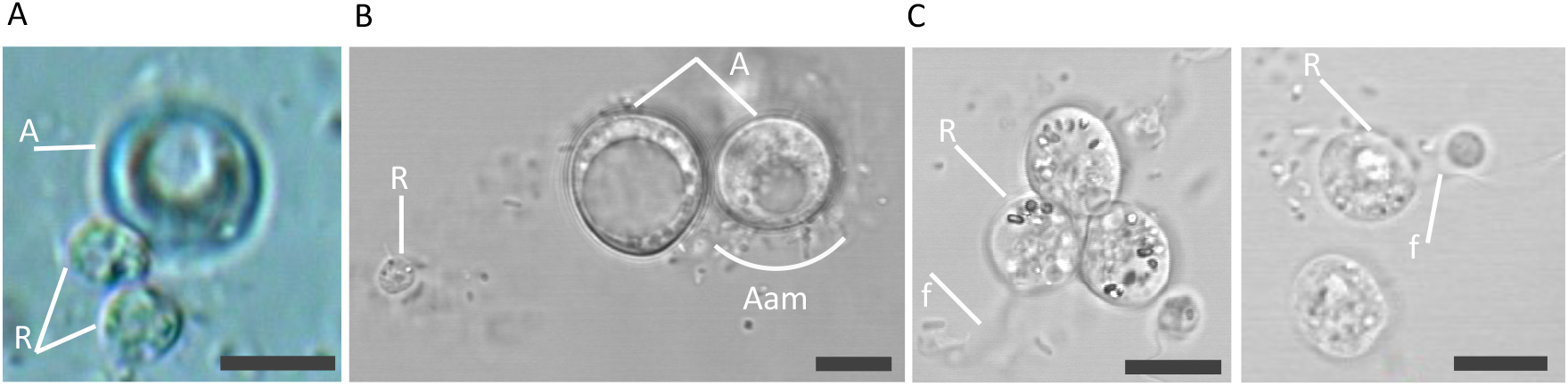
Possible interactions of *R. epiphylla* with *Albugo*. (A) Single cells of *Rhogostoma* (R) attached to the spore of *Albugo* (A). (B) *Albugo*’s spores associated with different microbes (Aam). (C) *Rhogostoma* cells feeding on other microbes or *Albugo*’s zoospores via filopodia (f) (See also S. Video1 and S. Video2). Measure bar indicates 10 μm.

**Supplementary Figure 7.**
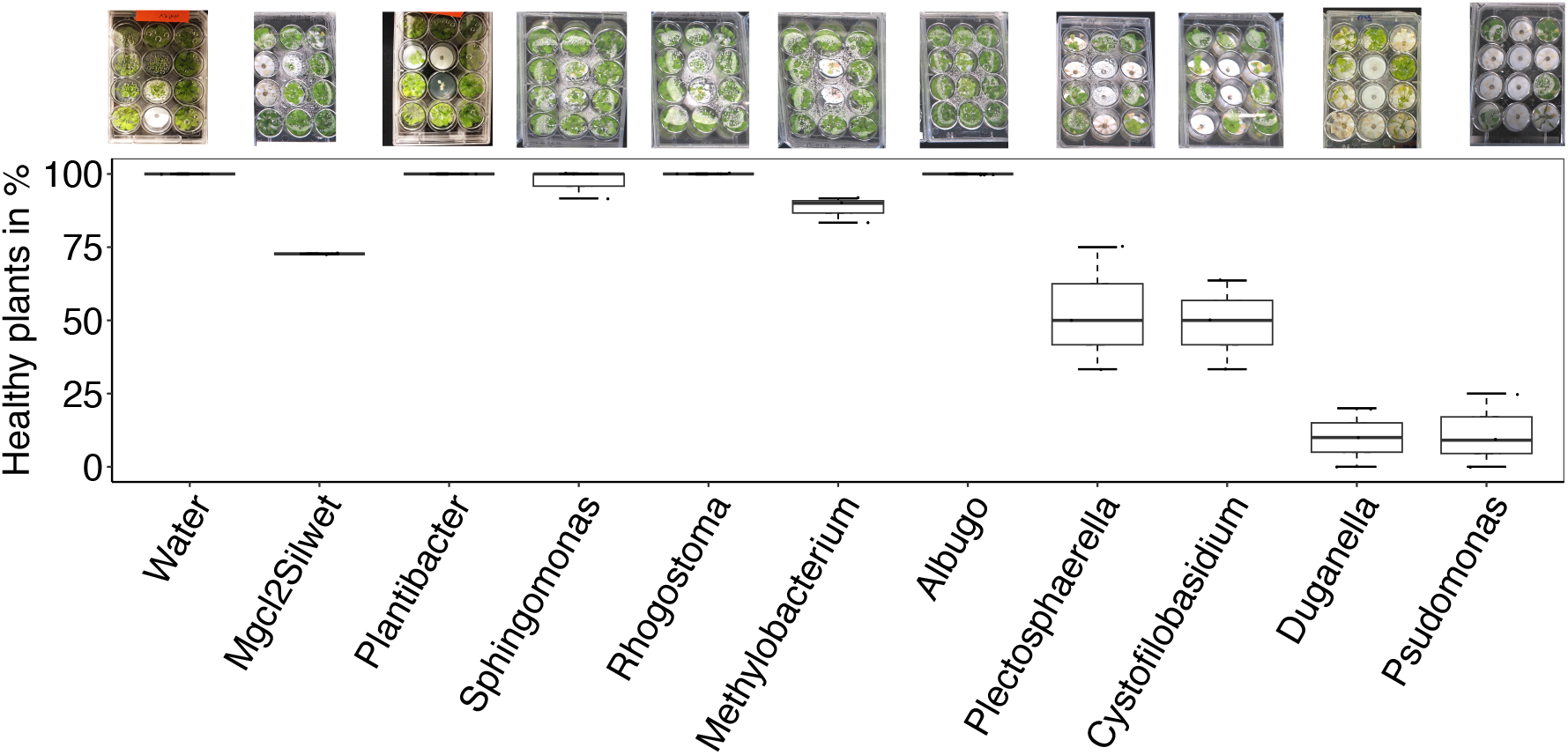
Effect of different HCom and Dcom microbes on WS-0 *Arabidopsis* plants under gnotobiotic conditions. Each microbe was sprayed on 3-4 weeks-old *Arabidopsis* plants under gnotobiotic conditions. Box plots show the percentage of healthy plants three weeks post-inoculation.

## Supplementary tables legends

**Table. S1. Detailing number of samples and sampling locations**.

**Table. S2. Results of blasting the sequences of key OTUs used in Figure 4 with NCBI databases**.

## Supplementary videos legends

**Supplementary Video S1 and S2. *Rhogostoma* grazing with their filopodia in the solution containing *Albugo*’s spores**.

## References

[1] Julia A Vorholt. Microbial life in the phyllosphere. Nature Reviews Microbiology, 10(12):828–840, 2012.

[2] Reza Sohrabi, Bradley C Paasch, Julian A Liber, and Sheng Yang He. Phyllosphere microbiome. Annual review of plant biology, 74:539–568, 2023.

[3] Brajesh K Singh, Manuel Delgado-Baquerizo, Eleonora Egidi, Emilio Guirado, Jan E Leach, Hongwei Liu, and Pankaj Trivedi. Climate change impacts on plant pathogens, food security and paths forward. Nature Reviews Microbiology, pages 1–17, 2023.

[4] Susan Trumbore, Paulo Brando, and Henrik Hartmann. Forest health and global change. Science, 349(6250):814–818, 2015.

[5] Anna Bonaterra, Esther Badosa, Núria Daranas, Jesús Franćes, Gemma Roselló, and Emilio Montesinos. Bacteria as biological control agents of plant diseases. Microorganisms, 10(9):1759, 2022.

[6] Sheridan L Woo, Rosa Hermosa, Matteo Lorito, and Enrique Monte. Trichoderma: a multipurpose, plant-beneficial microorganism for eco-sustainable agriculture, 2023.

[7] Zhenyan Zhang, Qi Zhang, Hengzheng Cui, Yan Li, Nuohan Xu, Tao Lu, Jian Chen, Josep Penuelas, Baolan Hu, and Haifeng Qian. Composition identification and functional verification of bacterial community in disease-suppressive soils by machine learning. Environmental Microbiology, 24(8):3405–3419, 2022.

[8] Michael J Sweet and Mark T Bulling. On the importance of the microbiome and pathobiome in coral health and disease. Frontiers in Marine Science, 4:9, 2017.

[9] Ricardo Hernández Medina, Svetlana Kutuzova, Knud Nor Nielsen, Joachim Johansen, Lars Hestbjerg Hansen, Mads Nielsen, and Simon Rasmussen. Machine learning and deep learning applications in microbiome research. ISME Communications, 2(1):98, 2022.

[10] Jun Yuan, Tao Wen, He Zhang, Mengli Zhao, C Ryan Penton, Linda S Thomashow, and Qirong Shen. Predicting disease occurrence with high accuracy based on soil macroecological patterns of fusarium wilt. The ISME Journal, 14(12):2936–2950, 2020.

[11] Hao-Xun Chang, James S Haudenshield, Charles R Bowen, and Glen L Hartman. Metagenome-wide association study and machine learning prediction of bulk soil microbiome and crop productivity. Frontiers in Microbiology, 8:519, 2017.

[12] Barbara Emmenegger, Julien Massoni, Christine M Pestalozzi, Miriam Bortfeld-Miller, Benjamin A Maier, and Julia A Vorholt. Identifying microbiota community patterns important for plant protection using synthetic communities and machine learning. Nature Communications, 14(1):7983, 2023.

[13] Eric Kemen, Anastasia Gardiner, Torsten Schultz-Larsen, Ariane C Kemen, Alexi L Balmuth, Alexandre Robert-Seilaniantz, Kate Bailey, Eric Holub, David J Studholme, Dan MacLean, et al. Gene gain and loss during evolution of obligate parasitism in the white rust pathogen of arabidopsis thaliana. PLoS biology, 9(7):e1001094, 2011.

[14] Juliana Almario, Maryam Mahmoudi, Samuel Kroll, Mathew Agler, Aleksandra Placzek, Alfredo Mari, and Eric Kemen. The leaf microbiome of arabidopsis displays reproducible dynamics and patterns throughout the growing season. Mbio, 13(3):e02825–21, 2022.

[15] Matthew T Agler, Jonas Ruhe, Samuel Kroll, Constanze Morhenn, Sang-Tae Kim, Detlef Weigel, and Eric M Kemen. Microbial hub taxa link host and abiotic factors to plant microbiome variation. PLoS biology, 14(1):e1002352, 2016.

[16] Daniel Gómez-Pérez, Monja Schmid, Vasvi Chaudhry, Yiheng Hu, Ana Velic, Boris Maček, Jonas Ruhe, Ariane Kemen, and Eric Kemen. Proteins released into the plant apoplast by the obligate parasitic protist albugo selectively repress phyllosphere-associated bacteria. New Phytologist, 2022.

[17] Maryam Mahmoudi, Juliana Almario, Katrina Lutap, Kay Nieselt, and Eric Kemen. Microbial communities living inside plant leaves or on the leaf surface are differently shaped by environmental cues. ISME Communications, 4 (1):ycae103, 08 2024. ISSN 2730-6151. doi: 10.1093/ismeco/ycae103. URL 10.1093/ismeco/ycae103.

[18] Sajjad Asaf, Muhammad Numan, Abdul Latif Khan, and Ahmed Al-Harrasi. Sphingomonas: from diversity and genomics to functional role in environmental remediation and plant growth. Critical Reviews in Biotechnology, 40(2):138–152, 2020.

[19] Lisa Röttjers and Karoline Faust. From hairballs to hypotheses–biological insights from microbial networks. FEMS microbiology reviews, 42(6):761–780, 2018.

[20] Christine M Vogel, Daniel B Potthoff, Martin Schäfer, Niculò Barandun and Julia A Vorholt. Protective role of the arabidopsis leaf microbiota against a bacterial pathogen. Nature microbiology, 6(12):1537–1548, 2021.

[21] Yong-Guan Zhu, Chao Xiong, Zhong Wei, Qing-Lin Chen, Bin Ma, Shu-Yi-Dan Zhou, Jiaqi Tan, Li-Mei Zhang, Hui-Ling Cui, and Gui-Lan Duan. Impacts of global change on the phyllosphere microbiome. New Phytologist, 2022.

[22] Nick C Snelders, Hanna Rovenich, Gabriella C Petti, Mercedes Rocafort, Grardy CM van den Berg, Julia A Vorholt, Jeroen R Mesters, Michael F Seidl, Reindert Nijland, and Bart PHJ Thomma. Microbiome manipulation by a soilborne fungal plant pathogen using effector proteins. Nature Plants, 6(11):1365– 1374, 2020.

[23] Katharina Eitzen, Priyamedha Sengupta, Samuel Kroll, Eric Kemen, and Gunther Doehlemann. A fungal member of the arabidopsis thaliana phyllosphere antagonizes albugo laibachii via a gh25 lysozyme. Elife, 10:e65306, 2021.

[24] Talia L Karasov, Manuela Neumann, Laura Leventhal, Efthymia Symeonidi, Gautam Shirsekar, Aubrey Hawks, Grey Monroe, Moisés Exposito-Alonso, Joy Bergelson, et al. Continental-scale associations of arabidopsis thaliana phyllosphere members with host genotype and drought. Nature Microbiology, pages 1–11, 2024.

[25] Wu Xiong, Rong Li, Yi Ren, Chen Liu, Qingyun Zhao, Huasong Wu, Alexandre Jousset, and Qirong Shen. Distinct roles for soil fungal and bacterial communities associated with the suppression of vanilla fusarium wilt disease. Soil Biology and Biochemistry, 107:198–207, 2017.

[26] Khaoula Belhaj, Liliana M Cano, David C Prince, Ariane Kemen, Kentaro Yoshida, Yasin F Dagdas, Graham J Etherington, Henk-jan Schoonbeek, H Peter van Esse, Jonathan DG Jones, et al. Arabidopsis late blight: infection of a nonhost plant by albugo laibachii enables full colonization by phytophthora infestans. Cellular Microbiology, 19(1):e12628, 2017.

[27] Dan Knights, Laura Wegener Parfrey, Jesse Zaneveld, Catherine Lozupone, and Rob Knight. Human-associated microbial signatures: examining their predictive value. Cell host & microbe, 10(4):292–296, 2011.

[28] Jens M Olesen, Jordi Bascompte, Yoko L Dupont, and Pedro Jordano. The modularity of pollination networks. Proceedings of the National Academy of Sciences, 104(50):19891–19896, 2007.

[29] Talia L Karasov, Juliana Almario, Claudia Friedemann, Wei Ding, Michael Giolai, Darren Heavens, Sonja Kersten, Derek S Lundberg, Manuela Neumann, Julian Regalado, et al. Arabidopsis thaliana and pseudomonas pathogens exhibit stable associations over evolutionary timescales. Cell host & microbe, 24(1):168–179, 2018.

[30] Or Shalev, Talia L Karasov, Derek S Lundberg, Haim Ashkenazy, Pratchaya Pramoj Na Ayutthaya, and Detlef Weigel. Commensal pseudomonas strains facilitate protective response against pathogens in the host plant. Nature ecology & evolution, 6(4):383–396, 2022.

[31] Derek S Lundberg, Roger de Pedro Jové, Pratchaya Pramoj Na Ayutthaya, Talia L Karasov, Or Shalev, Karin Poersch, Wei Ding, Anita Bollmann-Giolai, Ilja Bezrukov, and Detlef Weigel. Contrasting patterns of microbial dominance in the arabidopsis thaliana phyllosphere. Proceedings of the National Academy of Sciences, 119(52):e2211881119, 2022.

[32] Gerd Innerebner, Claudia Knief, and Julia A Vorholt. Protection of arabidopsis thaliana against leaf-pathogenic pseudomonas syringae by sphingomonas strains in a controlled model system. Applied and environmental microbiology, 77(10):3202–3210, 2011.

[33] Claudia Knief, Lisa Frances, and Julia A Vorholt. Competitiveness of diverse methylobacterium strains in the phyllosphere of arabidopsis thaliana and identification of representative models, including m. extorquens pa1. Microbial ecology, 60:440–452, 2010.

[34] Claudia Bartoli, Léa Frachon, Matthieu Barret, Mylène Rigal, Carine Huard-Chauveau, Baptiste Mayjonade, Catherine Zanchetta, Olivier Bouchez, Dominique Roby, Sébastien Carrère, et al. In situ relationships between microbiota and potential pathobiota in arabidopsis thaliana. The ISME journal, 12(8):2024–2038, 2018.

[35] María Florencia Gorordo, María Ester Lucca, and Marcela Paula Sangorrín. Biocontrol efficacy of the vishniacozyma victoriae in semi-commercial assays for the control of postharvest fungal diseases of organic pears. Current Microbiology, 79 (9):259, 2022.

[36] Maria Florencia Perez, Luciana Contreras, Nydia Mercedes Garnica, María Verónica Fernández-Zenoff, María Eugenia Farías, Milena Sepulveda, Jacqueline Ramallo, and Julián Rafael Dib. Native killer yeasts as biocontrol agents of postharvest fungal diseases in lemons. PloS one, 11(10):e0165590, 2016.

[37] Jia Liu, Michael Wisniewski, Samir Droby, Silvana Vero, Shiping Tian, and Vera Hershkovitz. Glycine betaine improves oxidative stress tolerance and biocontrol efficacy of the antagonistic yeast cystofilobasidium infirmominiatum. International Journal of Food Microbiology, 146(1):76–83, 2011.

[38] Kenneth Dumack, Sebastian Flues, Karoline Hermanns, and Michael Bonkowski. Rhogostomidae (cercozoa) from soils, roots and plant leaves (arabidopsis thaliana): Description of rhogostoma epiphylla sp. nov. and r. cylindrica sp. nov. European journal of protistology, 60:76–86, 2017.

[39] Michael Bonkowski. Protozoa and plant growth: the microbial loop in soil revisited. New Phytologist, 162(3):617–631, 2004.

[40] Kenneth Dumack, Christina Baumann, and Michael Bonkowski. A bowl with marbles: revision of the thecate amoeba genus lecythium (chlamydophryidae, tectofilosida, cercozoa, rhizaria) including a description of four new species and an identification key. Protist, 167(5):440–459, 2016.

[41] Bao-Anh Thi Nguyen, Kenneth Dumack, Pankaj Trivedi, Zahra Islam, and Hang-Wei Hu. Plant associated protists—untapped promising candidates for agrifood tools. Environmental Microbiology, 25(2):229–240, 2023.

[42] Sai Guo, Stefan Geisen, Yani Mo, Xinyue Yan, Ruoling Huang, Hongjun Liu, Zhilei Gao, Chengyuan Tao, Xuhui Deng, Wu Xiong, et al. Predatory protists impact plant performance by promoting plant growth-promoting rhizobacterial consortia. The ISME Journal, page wrae180, 2024.

[43] Nico Eisenhauer, Jes Hines, Fernando T Maestre, and Matthias C Rillig. Reconsidering functional redundancy in biodiversity research. npj Biodiversity, 2(1):9, 2023.

[44] Paul J McMurdie and Susan Holmes. phyloseq: an r package for reproducible interactive analysis and graphics of microbiome census data. PloS one, 8(4):e61217, 2013.

[45] Jari Oksanen, F Guillaume Blanchet, Roeland Kindt, Pierre Legendre, Peter R Minchin, R. O’hara, Gavin L Simpson, Peter Solymos, M Henry H Stevens, Helene Wagner, et al. Package ‘vegan’. Community ecology package, version, 2(9):1–295, 2013.

[46] Derek Ogle and Maintainer Derek Ogle. Package ‘fsa’. Cran Repos, 1:206, 2017.

[47] Core Team, Maintainer R Core Team, MASS Suggests, and S Matrix. Package stats. The R Stats Package, 2018.

[48] R R Core Team et al. R: A language and environment for statistical computing. 2013.

[49] Fabian Pedregosa, Gaël Varoquaux, Alexandre Gramfort, Vincent Michel, Bertrand Thirion, Olivier Grisel, Mathieu Blondel, Peter Prettenhofer, Ron Weiss, Vincent Dubourg, et al. Scikit-learn: Machine learning in python. the Journal of machine Learning research, 12:2825–2830, 2011.

[50] Jonathan Friedman and Eric J Alm. Inferring correlation networks from genomic survey data. PLoS computational biology, 8(9):e1002687, 2012.

[51] Stephen C Watts, Scott C Ritchie, Michael Inouye, and Kathryn E Holt. Fastspar: rapid and scalable correlation estimation for compositional data. Bioinformatics, 35(6):1064–1066, 2019.

[52] Aric Hagberg, Pieter J Swart, and Daniel A Schult. Exploring network structure, dynamics, and function using networkx. Technical report, Los Alamos National Laboratory (LANL), Los Alamos, NM (United States), 2008.

[53] Vincent D Blondel, Jean-Loup Guillaume, Renaud Lambiotte, and Etienne Lefebvre. Fast unfolding of communities in large networks. Journal of statistical mechanics: theory and experiment, 2008(10):P10008, 2008.

[54] Paul Shannon, Andrew Markiel, Owen Ozier, Nitin S Baliga, Jonathan T Wang, Daniel Ramage, Nada Amin, Benno Schwikowski, and Trey Ideker. Cytoscape: a software environment for integrated models of biomolecular interaction networks. Genome research, 13(11):2498–2504, 2003.

